# TaxTriage: An Open-Source Metagenomic Sequencing Data Analysis Pipeline Enabling Putative Pathogen Detection

**DOI:** 10.1101/2025.07.16.664785

**Authors:** Brian Merritt, Jeremy D Ratcliff, Stanley Ta, Gunars Osis, Matthew R. Mauldin, Peter M Thielen

## Abstract

**Motivation:** TaxTriage is a comprehensive pathogen identification workflow designed for both short– and long-read untargeted DNA and RNA sequencing data. Combining read classification, mapping, and *de novo* assembly approaches, putative pathogens are identified through comparisons to curated pathogens and abundance expectations from healthy cohort data. Flexible installation options are enabled using Nextflow™ (NF), including cloud deployment via NF Tower (Seqera Platform) and local installation on a variety of systems, including standalone installations without external internet access. Final analysis summaries are compiled into an Organism Discovery Report, which lists likely pathogens and supporting data, including a custom confidence score.

**Results:** Evaluation of published *in silico*, clinical, and outbreak datasets identified performance comparable to alternative cloud-based processing pipelines for expected pathogen and co-infection detection with similar sensitivity and increased specificity. To support both public health and veterinary diagnostics communities, customization options have been incorporated to enable improved performance for host species of interest.

**Availability and Implementation:** Source code for TaxTriage is freely available at https://github.com/jhuapl-bio/taxtriage.

## Introduction

The increased adoption of next generation sequencing (NGS) instruments in diagnostic laboratories has dramatically improved pathogen characterization capabilities ^1–3^. Sequencing of total DNA or RNA from a clinical sample has shown great promise for untargeted pathogen detection, having been deployed in highly capable clinical laboratories to great effect ^4–8^. Widespread adoption of genomics technologies for pathogen identification and characterization remains limited due to several factors, including cost, regulatory uncertainty, data management, and complex data analysis processes that require specialized training to interpret analysis outputs^9–12^.

Metagenomic applications can utilize samples that have either been enriched for specific sequences (targeted) or take an appraisal of total input nucleic acids (untargeted). Techniques for targeted approaches include physical host depletion, targeted amplification, and hybrid capture enrichment based on oligonucleotide probes ^13–15^. While targeted sequence capture panels can increase the sensitivity for specific, known nucleic acid sequences ^16^, untargeted metagenomic next-generation sequencing (mNGS) provides a less-biased representation of microbial communities, increassing the taxonomic breadth of organisms that can be simultaneously detected. For example, untargeted mNGS was used to produce several of the first sequences of SARS-CoV-2 in late 2019 ^17^. mNGS has been increasingly paired with diagnostic assays at the outset of new outbreaks to evaluate phylogenetic diversity, as demonstrated mid-2022 with the spread of mpox ^18^. Untargeted mNGS of complex samples can also present unique challenges, such as higher limits of detection due to the untargeted nature, as well as the preponderance of uninformative host or other background nucleic acids within a clinical sample, which reduce the proportion of microbial data available to identify a pathogen.

Within public health and clinical laboratory settings, major barriers to mNGS adoption include the computational infrastructure required to generate, house, and analyze large data files, as well as the technical expertise for interpretation of analytical outputs. Typical laptop computers lack sufficient processing power to perform mNGS analyses, and interacting with outputs of bioinformatic algorithms frequently requires engaging with data from a command line interface. In addition, deriving public health insights from mNGS data typically requires a nuanced understanding of the outputs. These requirements limit the implementation of mNGS following distributed placement of NGS platforms, due to budget constraints, technical expertise, and turnaround times.

Here, we present TaxTriage, a fully open-source informatics pipeline for untargeted mNGS data analysis. TaxTriage enables end-to-end processing of raw sequencing data, generated from short-or long-read platforms, to produce species-level identification of potential pathogens. The underlying pipeline limits user interaction with intermediate files, producing an Organism Discovery Report that summarizes derived findings from intermediate mNGS analytical outputs. Default settings produce a confidence metric for each detected organism, and optional modules improve identification of known or suspected human pathogens. Flexible software deployment options enable local processing of human clinical samples, containing potentially sensitive genetic information, without requiring third-party processing services or the internet. By lowering the barrier to entry for mNGS data analysis, TaxTriage democratizes access to threat agnostic sentinel surveillance capabilities using existing NGS instruments in public health and clinical laboratories. This manuscript describes development through v2.0.8 (https://github.com/jhuapl-bio/taxtriage/releases/tag/v2.0.8).

## System, methods, and usage

TaxTriage is an end-to-end bioinformatics pipeline developed using the command line Nextflow™ (NF) computational workflow ^19^, coupling support from the nf-core platform for implementing a variety of informatics libraries and modules ^20^. By leveraging nf-core, all data processing steps are performed consecutively without expert input or the manual conversion of file types for downstream processing. TaxTriage is provided as open-source software (OSS) and can be deployed using either Docker or Singularity containers, enabling use in local, high-performance computing (HPC), and cloud-based environments (e.g., Amazon Web Services [AWS], Google Cloud, and Microsoft Azure). While the default pipeline requires an internet connection to download reference sequences from public repositories, TaxTriage can be configured to use a locally held sequence database and run in an offline or air-gapped computational environment. TaxTriage source code and documentation are available on GitHub: https://github.com/jhuapl-bio/taxtriage.

TaxTriage lowers the barriers required to implement metagenomic sequencing in clinical and public health laboratories. To automate the processing of sequence reads, TaxTriage chains together standard, publicly available bioinformatic utilities that would typically be implemented by an mNGS bioinformatics specialist (Figure 1). These processes cover all steps from quality control (QC), *in silico* host sequence removal, metagenomic classification, alignment, and assembly, ultimately culminating in a single output—a portable document format (PDF) Organism Discovery Report (ODR). The ODR lists organisms identified within mNGS datasets, along with supporting metadata and a unified confidence metric (TASS Score), to increase the report’s utility for clinical and public health professionals. The pipeline uses a variety of custom-developed and publicly supported nf-core modules (Table 1). There is flexibility in the execution of TaxTriage, with many modules enabled by default (“Primary”), and custom combinations of primary and supplemental modules available based on user input.

**Figure 1:**
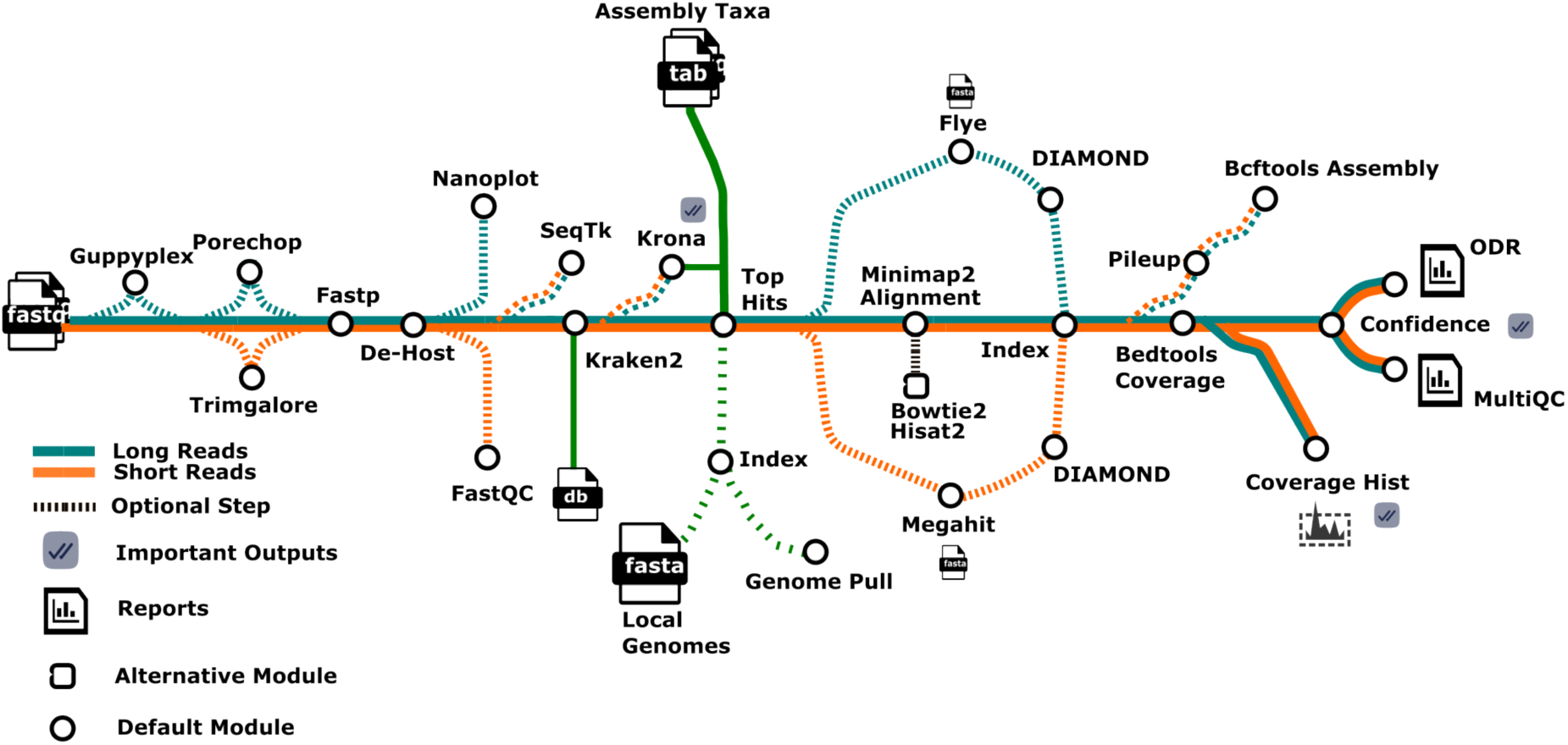
Workflow diagram of each step for both Illumina and Nanopore data analysis. Each module is called sequentially to create the final MultiQC and Organism Discovery Report. Some steps are optional to reduce overall computational complexity and are defined through calls within the NF platform. Broadly, “Nanopore” tracks can be used for long-read sequencing data while “Illumina” tracks can be used with short-read sequencing data. Consistent with TaxTriage release v2.0.8.

**Table 1.** Bioinformatic processes utilized by TaxTriage.

| Process<br>(In Operation Order) | Module(s) | Primary /<br>Supplemental |
| --- | --- | --- |
| Read Preprocessing | Guppyplex (long-read) | Supplemental |
| Trimming | Trimgalore (short-read) <sup>21</sup>, Porechop (long-read) <sup>22</sup> | Supplemental |
| Quality Control (QC) | FASTP (Both) <sup>23</sup> | Primary |
| Read Normalization | Khmer (Both) <sup>24</sup> | Supplemental |
| Host(s) Removal | Minimap2 (Both) <sup>25</sup> | Primary |
| QC Plotting | FASTQC (short-read) <sup>26</sup>, Nanoplot (long-read) <sup>27</sup> | Primary |
| Metagenomic | Kraken2 (Both) <sup>28</sup> | Primary |
| Abundance Visualization | Krona (Both) <sup>29</sup> | Primary |
| Core Hits Filtering | Custom (Both) | Primary |
| Reference Retrieval | Custom (Both) | Primary |
| Nucleotide Alignment | Minimap2 (Both) <sup>25</sup>, Bowtie2 (short-read) <sup>30</sup>, Hisat2 | Primary |
| Protein Alignment | DIAMOND (Both) <sup>32</sup> | Supplemental |
| Alignment Statistics | Samtools (Both) <sup>33</sup> | Primary |
| Reference Assembly | Bcftools (Both) <sup>34</sup> | Supplemental |
| *De novo* Assembly | MEGAHIT (short-read) <sup>35</sup>, Flye (long-read) <sup>36</sup> | Supplemental |
| Pathogen Report | Custom + ReportLab <sup>37</sup> (Both) | Primary |
| MultiQC Report | MultiQC (Both) <sup>38</sup> | Primary |

Bioinformatic processes utilized by TaxTriage (Table 1) are enabled as “Primary” by default. Each module can be disabled or skipped based on additional optional parameters. “Supplemental” indicates that the process is optional and is disabled by default. Each custom-developed module leverages a variety of Python, bash, and/or NF-derived processes. The selection of a nf-core tool for each step is based on read length (e.g., short-or long-read chemistries) for each sample and is reported in the Module(s) column. Short-read modules have been tested with data generated by Illumina, Pacific Biosciences (PacBio), and Element Biosciences instruments; long-read modules have been leveraged with data originating from Oxford Nanopore Technologies (ONT) and PacBio instruments.

The primary TaxTriage pipeline requires raw NGS reads in compressed or uncompressed.fastq format and a sample sheet describing the input data in Comma-Separated Values (CSV).csv format. A sample sheet requires metadata for sample names, sequencing platform used, and.fastq file paths, and can optionally include body site value (one of stool, nasal, oral, skin, or vaginal). After a series of validation checks for accurate designation of the NGS platform, read path(s), and naming convention, there is an optional step for read trimming using either Porechop or Trimgalore for long-read or short-read data, respectively. Fastp filters low-quality reads while FastQC (short-read) and Nanoplot (long-read) generate supplementary QC plots. Host DNA is removed by minimap2 (disabled by default) at this stage, targeting a user-specified species that is provided either as a genome code or as a local fasta file. Cleaned reads are classified using Kraken2, although additional classifiers are actively being incorporated. For Kraken2-based classification, users can either upload a custom Kraken2 database or leverage NF support to pull a variety of default databases defined and provided publicly on Kraken2’s AWS indices site (https://benlangmead.github.io/aws-indexes/k2).

Minimap2 (long-reads) and Bowtie2 (short-reads) are both available for nucleotide sequence alignment; protein alignment of de novo assembled contigs can be optionally performed using DIAMOND. The accuracy, sensitivity, and computational cost of sequence alignment are heavily dependent on the size and completeness of the reference database used. In TaxTriage with access to the external internet, the default pipeline uses the classification output to identify the species to be downloaded from the National Center for Biotechnology Information (NCBI) file transfer protocol (FTP) genomes repository of RefSeq representative assemblies. Users can change the total number of organism assemblies to download; by default, this value is the 400 species with the highest read abundance based on Kraken2 classification. TaxTriage includes multiple mechanisms for enhancement of alignment-based pathogen detection, such as use of a curated pathogen list containing over 1700 targets, which can be run in conjunction with abundance-based download limits set by the user. Alternatively, a local alignment database can be used if particular organisms are of interest, or the deployment does not have access to the external internet. Contig assembly is also available as an optional module and is required for DIAMOND.

TaxTriage produces a summarized output PDF that consolidates alignment statistics and confidence metrics for each identified species (“Organism Discovery Report”; Figure 2). Below, we describe the calculation and definition of columns in the Organism Discovery Report:

**Figure 2:**
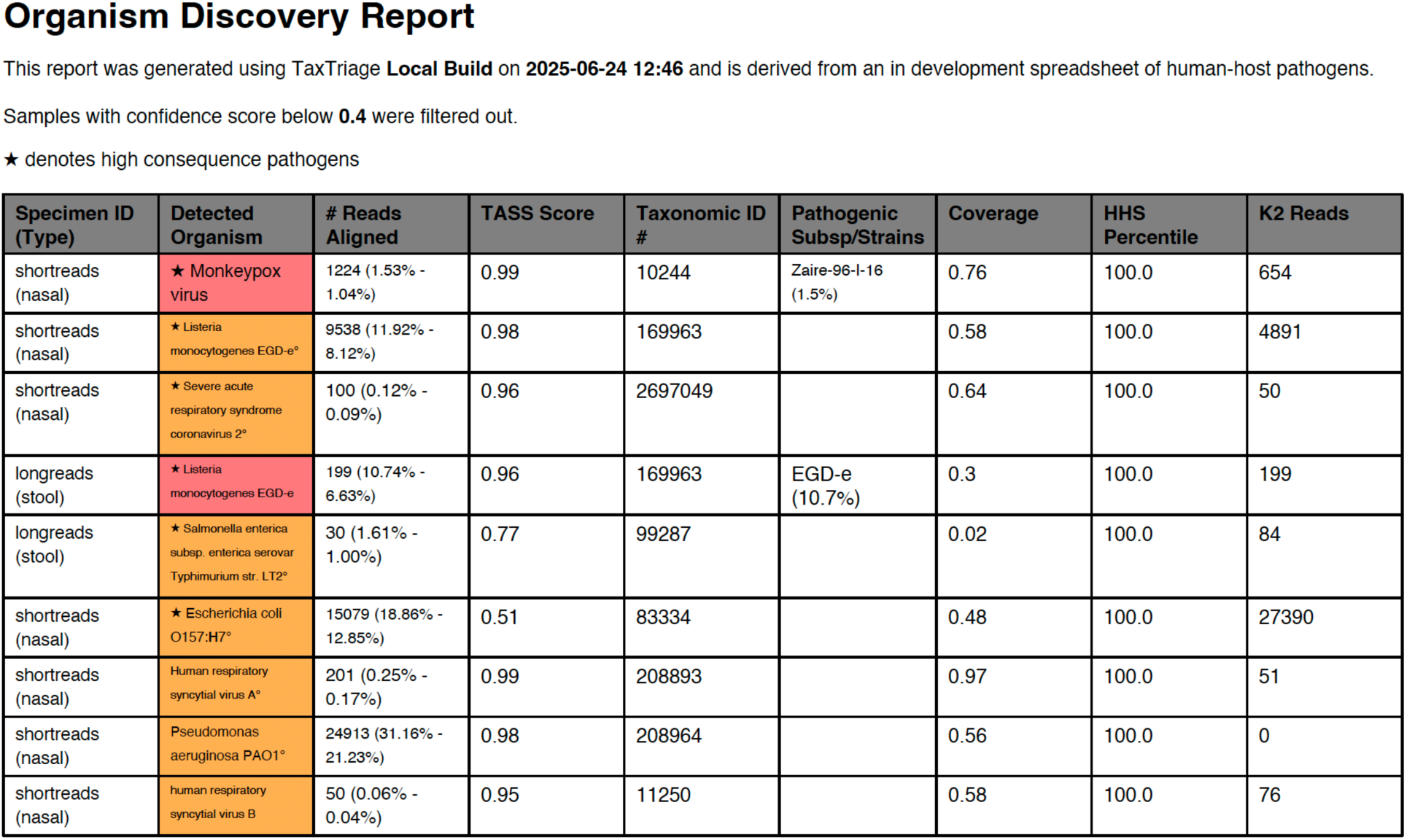
Example Organism Discovery Report Output. General alignment statistics (including read counts) are reported at the start of the report for all defined pathogens and commensals (disabled by default; displayed in lower table). Those marked with a star denote high-consequence pathogens, including those on the NIAID biodefense pathogen list ^39^. Organisms without class labels (i.e., not categorized/classified as pathogenic or commensal) are reported in the supplementary Unannotated Organisms list. This example report was generated from multiple *in silico* data samples, generated with various depth of coverage profiles using InSilicoSeq and references were curated from NCBI’s Refseq database ^40^. Additional annotation explanations are available later in the report (not depicted in the sample figure). Supplemental or intermediate files are available in output directories according to their function within the workflow and can be used according to additional bioinformatics downstream requirements if needed.

### Organism Detection and Confidence

A core metric describing organism detection is the raw value of all reads that have aligned to a reference genome with a configurable map quality (MapQ) score of at least 20 (default). Species detection is defined as at least three non-overlapping reads aligning to a reference genome with MapQ scores above the configured threshold. One challenge with organism detection via metagenomics is the risk of inaccurate reporting of organism presence, particularly erroneous reporting of high-consequence pathogens or select agents. In order to reduce the risk of false positives, all detected organisms have an accompanying confidence score (“TASS Score”). This unitless metric that ranges from 0 to 1 combines multiple approaches to measure genome coverage inequality, which, from the experience of our team and conversations with the community, can be a representation of improper nucleotide alignments (i.e., multiple reads only aligning to specific regions that are conserved in closely related organisms). The TASS Score is generated through an Average Fair Distribution Score (*AFDS*; consideration of inequality of coverage and depth), Breadth of Coverage Score (*BoCS*; proportion of reference genome coverage represented by input data), and Comparison Hash Score (*CHS*; MinHash sketches capturing unique k-mer content).

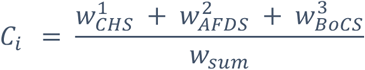

**Equation 1: TASS Score Generation.** Weighted scoring is applied to generate a value between 0-1, indicating overall confidence in pathogen presence. A global optimization leveraging *in silico* generated data established weights for Comparison Hash Score (*w^1^CHS*=5%), Average Fair Distribution (*w^2^AFDS*=71%), and Breadth of Coverage (*w^3^BoCS*=24%). Additional information on individual TASS Score components can be found in the Supplementary Materials.

By default, only pathogens with TASS scores at or above a threshold of 0.4 are listed in the Organism Discovery Report; this threshold can be user-configured as needed. The only exceptions are for organisms on the NIAID list of Biodefense Pathogens, which are tagged as high-consequence and listed in a separate table regardless of TASS score (Figure S1).

### Pathogen Classification

To enhance public health utility, each microorganism listed within the Organism Discovery Report is categorized according to standard, supported pathogen classifications: commensal, opportunistic, potential, or primary (Table 2). These species-level labels are derived from matching organism taxonomy IDs with a curated pathogen annotation sheet, which is stored as.csv and .txt files on the TaxTriage GitHub repository. This datasheet contains aggregate results from other efforts to document known human pathogens, namely Bartlett et al. (2022) ^41^, Taylor et al. (2001) ^42^, the WHO fungal priority pathogens list ^43^, the NIAID list of Biodefense Pathogens ^39^, the HHS and USDA Select Agents and Toxins List ^44^, and the CZID pathogens list ^45^. The pathogen annotation table includes organism name, NCBI taxonomy IDs, pathogen classification, known commensal and pathogenic body sites, and supportive references. These annotations can be further customized for additional body sites, host species, or alternative applications.

**Table 2.**
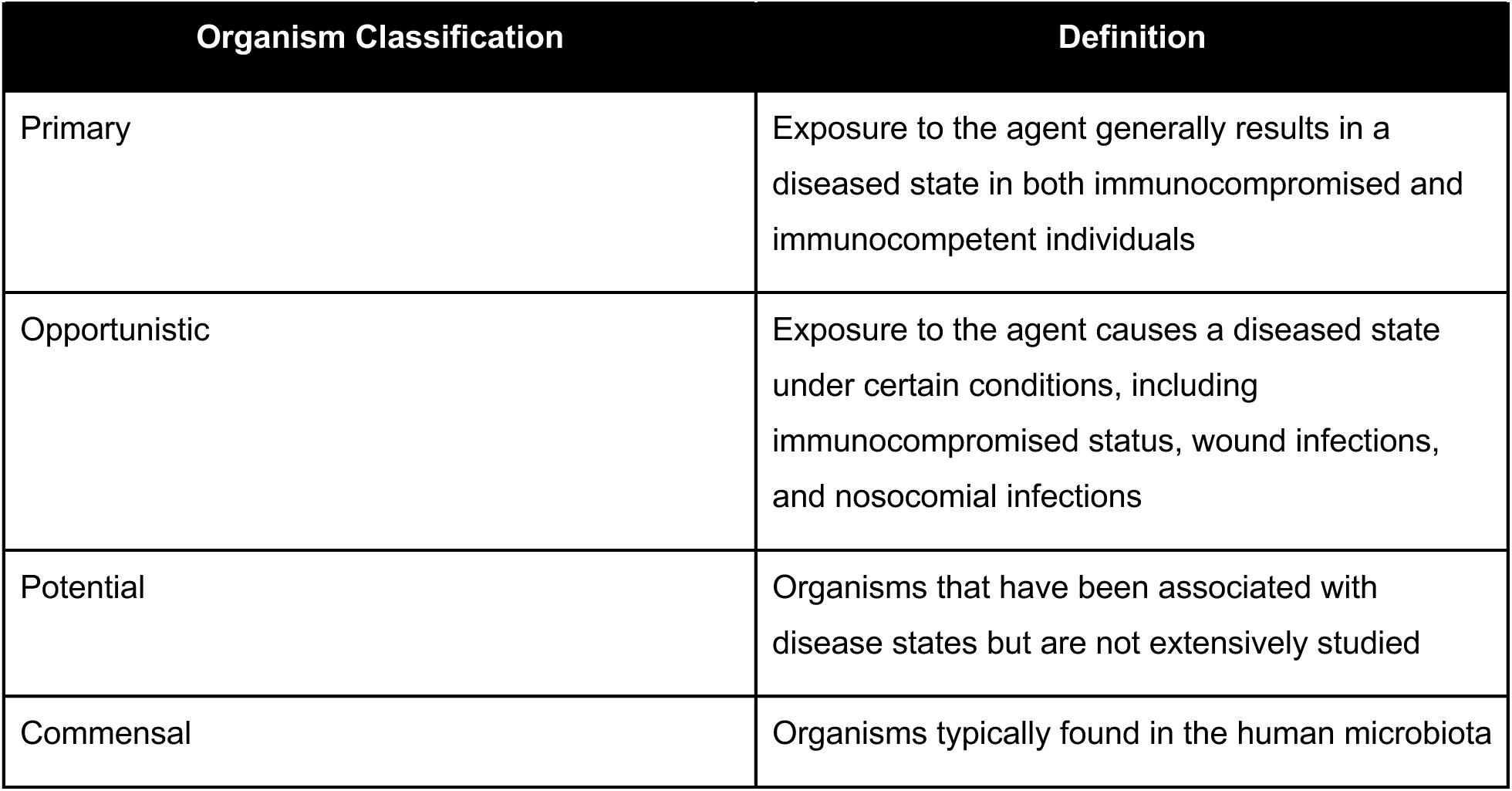
Organism Classifications used in Pathogen Annotation Table.

| Organism Classification | Definition |
| --- | --- |
| Primary | Exposure to the agent generally results in a diseased state in both immunocompromised and immunocompetent individuals |
| Opportunistic | Exposure to the agent causes a diseased state under certain conditions, including immunocompromised status, wound infections, and nosocomial infections |
| Potential | Organisms that have been associated with disease states but are not extensively studied |
| Commensal | Organisms typically found in the human microbiota |

For organisms classified as suspected pathogens when found in particular body sites, the pathogen will only be flagged if the sample originates from a matched body site as included in the input sample sheet. If the sample collection site is not specified, analysis proceeds without considering body site-specific pathogen annotation. In most cases, defining these assignments required targeted literature searching; the publications supporting the sample site designations are provided in the curated pathogen annotation table. In addition to providing organism metadata in the final report, a custom alignment database composed of species present in the pathogen annotation sheet can be used to supplement the top 400 (default) organisms identified via classification, bypassing the need for organism identification via classification for alignment and increasing sensitivity for known pathogens.

We recognize this list of pathogens is non-exhaustive and will require frequent curation. By storing the reference sheet on GitHub, members of the community can open issues or submit pull requests to add additional organisms via an approval system, increasing sensitivity for all users. They may also modify locally held versions of the sheet prior to launching a job, enabling a focus on organisms of particular interest to the user.

### Relative Abundance in Healthy Human Subjects

While the pathogen sheet increases sensitivity for known, curated pathogens, it is not adequate for identifying organisms absent from the database. TaxTriage contains an additional mechanism to identify abnormal distributions of commensal organisms or organisms not known to typically infect humans. Datasets from the NIH Human Microbiome Project (HMP) consist of over 108,000 whole-genome sequencing runs on ILL platforms derived from publicly available healthy human subjects (HHS) submitted to the sequence read archive (SRA) as of May 7, 2024 ^46^. From this data, expected relative distributions were generated per organism per body site (see figure 3 for sample breakdown between body sites). Within a TaxTriage sequencing run, the relative abundance of each organism is calculated. By default, an organism is flagged for follow-up if the observed abundance differs from the expected distribution by a z-score of >= 1.5. Additionally, organisms that have never been observed in the HMP datasets are flagged. Results from Kraken2, pathogen labels, and abundance calculations influence the content and organization of the Organism Discovery Report (Figure 4). The identification of commensal at “normal” abundance levels also reduces the total number of organisms present in the Organism Discovery Report, focusing users on the high likelihood causes of pathology.

**Figure 3:**
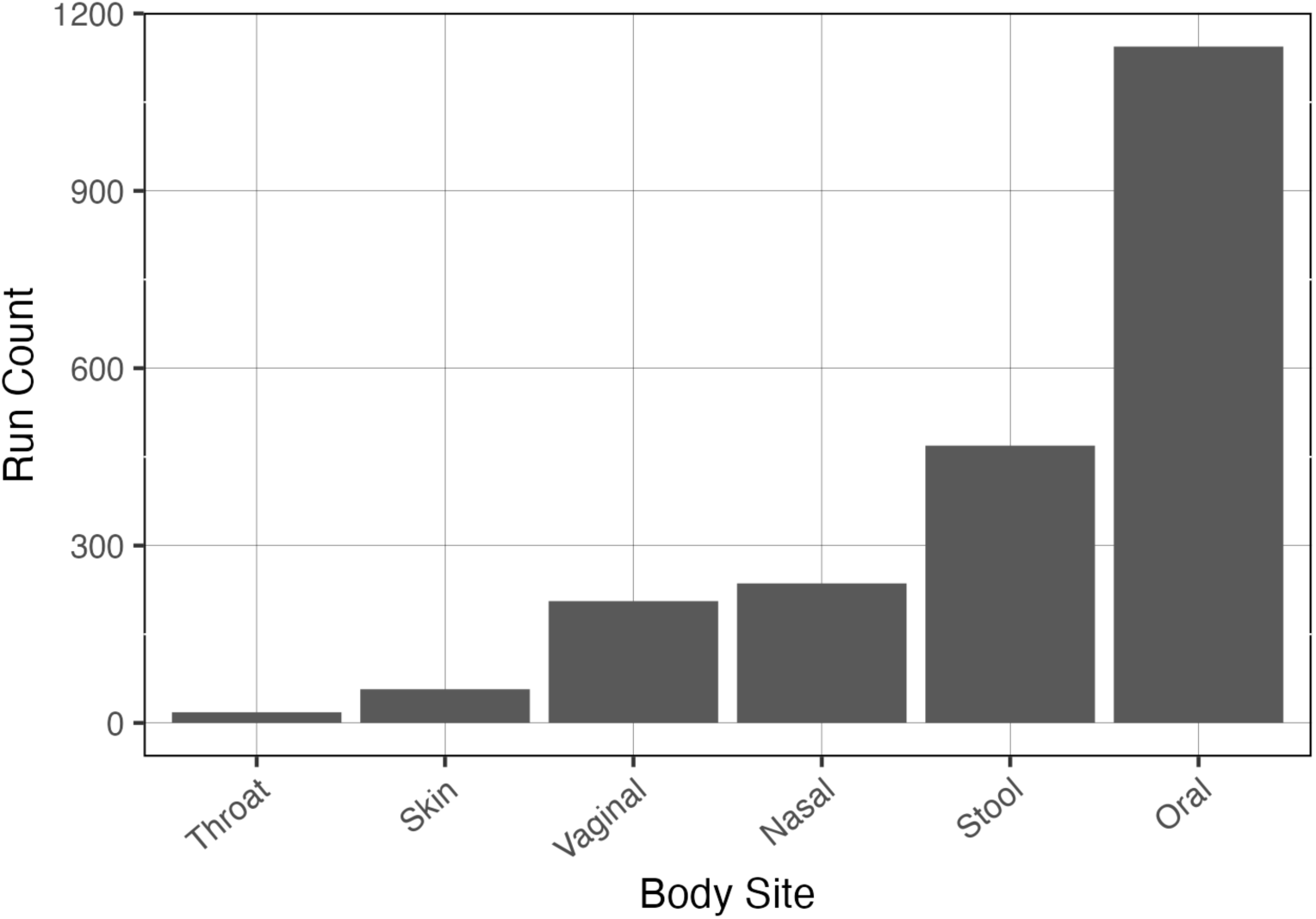
Count of run types based on body site for the HHS subjects. Generally, sequencing runs do not equate to the total number of SRA submissions (108k) available in the entire set of publicly available HHS datasets. Body sites with a greater number of sample runs will provide more precise relative abundance values for the irregular abundance derivation. Data as March 7, 2024.

**Figure 4:**
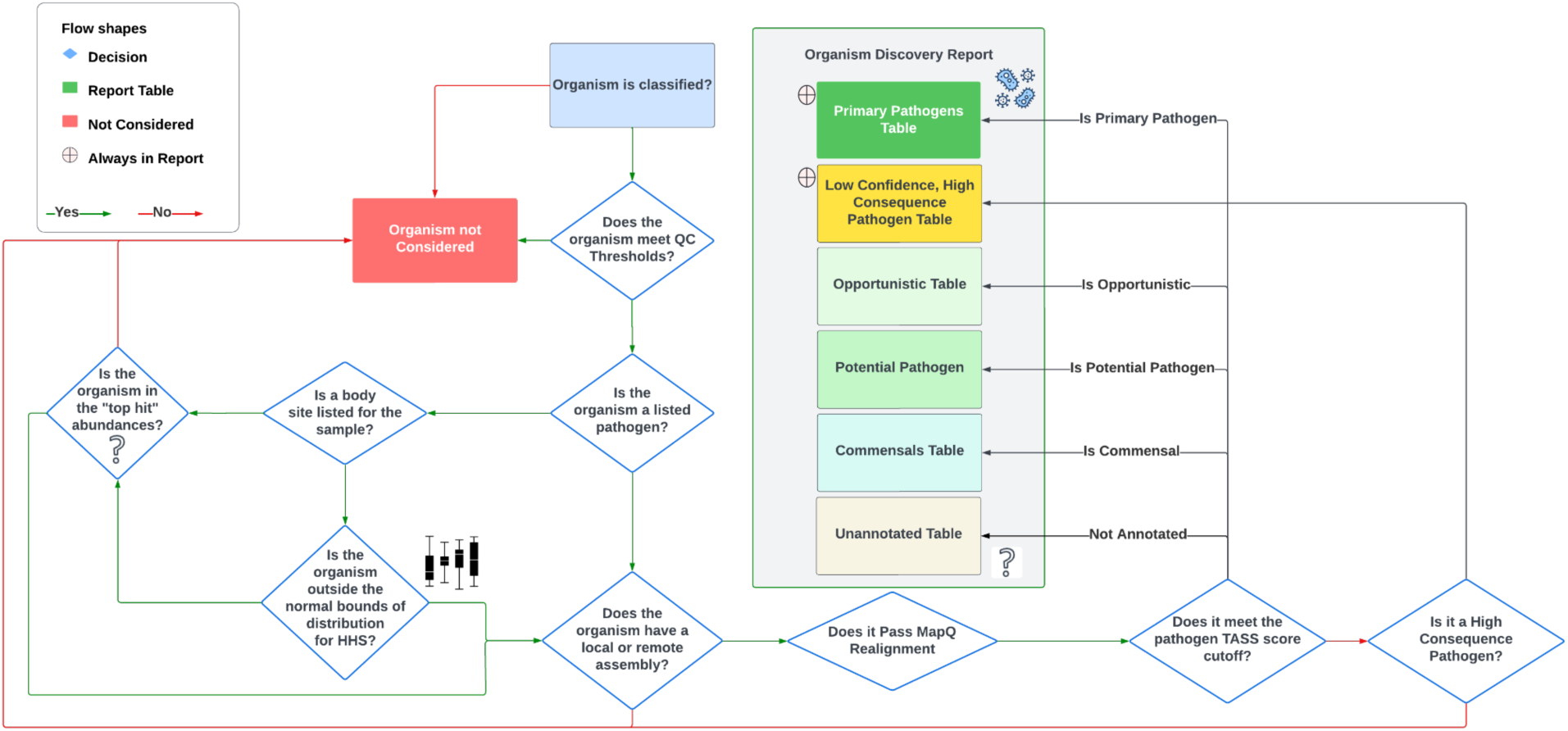
Decision tree for down-sampling organism classifications at various steps for assignment within the Organism Discovery Report. Proper placement of an organism within downstream steps and its correct assignment is based on multiple factors. Highest priority taxa will appear at the top, sorted on highest relative abundance in the given sample(s). This method heavily relies on specifying a supported sample type (body site) such as nasal, stool, or blood. HMP data is incorporated to ensure adequate removal of anticipated commensals to further distill the report and ease interpretation by focusing on microbes with higher likelihood of health implications. Presence of commensal, opportunistic, potential, or unannotated organisms are defined using workflow-specific flags.

### Additional Features and Ongoing Improvements

In addition to the core functionality, there are several supplementary features already incorporated or in development for TaxTriage. This includes non-human host identification and filtering, automated analysis of coding regions (CDs), and incorporation of *in silico* positive controls. Also, a built-in test dataset to analyze is required to complete without error prior to a version release. As TaxTriage is OSS under continuous development, additional features will continue to be incorporated based on feedback from the community. GitHub’s branching feature allows users operating under quality management systems to use stable versions.

## Benchmarking TaxTriage

To benchmark TaxTriage performance with other publicly available and custom-built analysis pipelines, we executed the pipeline on an *in silico* dataset, a “real-world” clinical dataset, and a “real-world” outbreak dataset. The *in silico* dataset enables comparison against ground truth organism presence; the clinical and outbreak datasets are an appropriate simulation of TaxTriage performance in an operational setting. As a common comparator, we additionally processed the datasets using the web GUI from the Chan Zuckerberg Initiative (CZID), using the Illumina pipeline version 8.3 aligned against the NCBI Nucleotide (NT)/non-redundant protein (NR) database indexed on 2024.02.06.

The *in silico* dataset is composed of two single-end “runs” generated as part of the COMPARE (Collaborative Management Platform for Detection and Analyses of (Re-)emerging and Foodborne Outbreaks in Europe) program ^47^. The datasets contain over 6 million simulated reads from a 150-bp Illumina HiSeq 2500 system (Table 3) and include four pathogenic viruses spiked in at varying depths and coverages within a background where 99.9% of the reads were derived from human or commensal bacteria. This dataset afforded two noted advantages: (1) the COMPARE publication provides rich information on the performance of 13 diverse, custom analysis pipelines against which TaxTriage can be compared; and (2) the COMPARE dataset includes reads derived from a “novel” avian bornavirus which enable a stress-test of the utility of TaxTriage to identify novel pathogens.

**Table 3.**
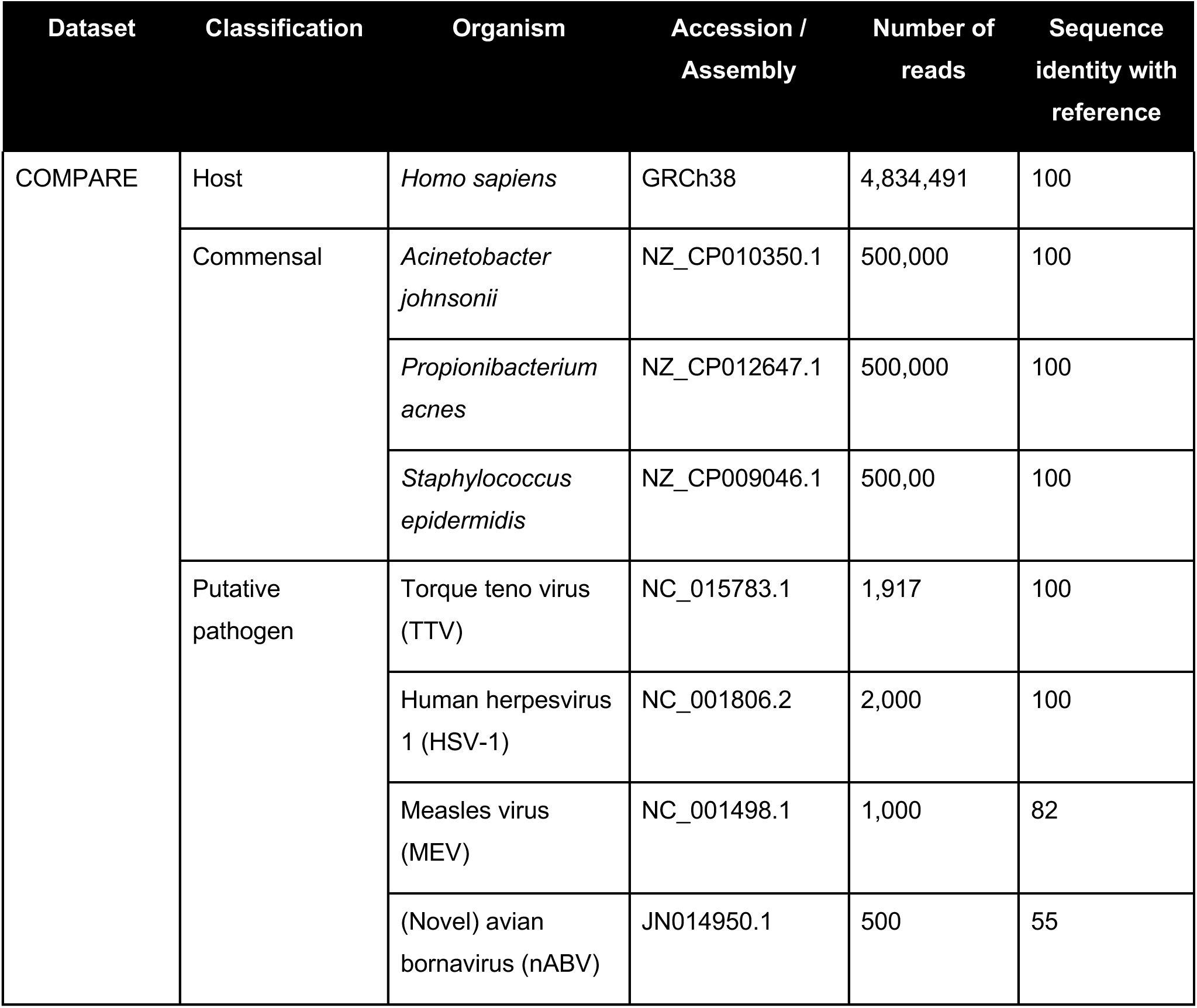
Characteristics of *in silico* comparison dataset.

| Dataset | Classification | Organism | Accession / Assembly | Number of reads | Sequence identity with reference |
| --- | --- | --- | --- | --- | --- |
| COMPARE | Host | *Homo sapiens* | GRCh38 | 4,834,491 | 100 |
|  | Commensal | *Acinetobacter johnsonii* | NZ_CP010350.1 | 500,000 | 100 |
|  |  | *Propionibacterium acnes* | NZ_CP012647.1 | 500,000 | 100 |
|  |  | *Staphylococcus epidermidis* | NZ_CP009046.1 | 500,00 | 100 |
|  | Putative pathogen | Torque teno virus (TTV) | NC_015783.1 | 1,917 | 100 |
|  |  | Human herpesvirus 1 (HSV-1) | NC_001806.2 | 2,000 | 100 |
|  |  | Measles virus (MEV) | NC_001498.1 | 1,000 | 82 |
|  |  | (Novel) avian bornavirus (nABV) | JN014950.1 | 500 | 55 |

The clinical dataset comes from the Initiative for the Critical Assessment of Metagenome Interpretation (CAMI) II clinical pathogen detection challenge ^48^. The dataset contains 6.9 million 100-bp reads from a paired-end run on an Illumina MiSeq. Host reads have been removed and replaced by the challenge organizers to protect anonymity of the patient. The challenge instructed participants to identify a causative agent of infection based on the metagenomic sequencing results and a case description. While the project metadata does not provide ground truth read counts, the challenge publication does provide the results from n = 10 community submissions. The target pathogen was Crimean-Congo Hemorrhagic Fever Virus (CCHFV).

Lastly, the real-world dataset is composed of sequencing data generated during the 2014-2016 West African Ebola outbreak in *Gire et al. (2014)* ^49^. Raw reads generated from Illumina sequencing of blood samples from n = 20 randomly selected patient samples suspected of Ebola Virus Disease (EVD) were downloaded from the Sequence Read Archive (SRA). Pathogens of interest were Zaire ebolavirus and any additional coinfections.

Overall results from pathogen identification in the test datasets are reported in Table 4. In general, TaxTriage successfully identified the target pathogens (COMPARE, CAMI) or likely causative agents of disease (*Gire et al. (2014)*). The exceptions were the mutated MEV and nABV targets in the COMPARE study. While MEV was detected at low levels in one of the runs, it fell below the 0.4 threshold for pathogen identification. In contrast, CZID identified substantially more MEV reads than TaxTriage. In fact, across nearly all sample-pathogen pairs, CZID reports a higher number of aligned nucleotide reads. This difference is expected as CZID aligns against the entirety of NCBI’s nucleotide (NT) database versus the single reference sequence used per species in TaxTriage. This difference in approach enables CZID to more readily identify organisms distinct from reference (provided there is representation within NCBI), but at the cost of reduced specificity and increased computational load. There are also likely other differences in alignment cutoffs between the two pipelines, as only 62 reads mapped with MapQ scores > 20 to a custom alignment database of all 23,533 MEV genomes on NCBI, far fewer than the 958 identified by CZID. Read recovery rates were, as expected, higher for CZID than TaxTriage on this dataset (Figure S2).

**Table 4.** Pathogens identified by TaxTriage in benchmarking datasets.

| Study | Sample/run | Pathogen | TASS Score | TaxTriage Mapped Reads | CZID Mapped Reads (NT) | CZID Mapped Reads (NR) |
| --- | --- | --- | --- | --- | --- | --- |
| CAMI | patmg | CCHFV | 0.50 | 22 | 177 | 90 |
|  |  | HIV-1 | 0.51 | 28 | 40 | 40 |
| COMPARE | ERR3313866 | TTV | ND | ND | 939 | 939 |
|  |  | HSV-1 | 0.95 | 756 | 841 | 818 |
|  |  | MEV | 0.22 | 3 | 465 | 489 |
|  |  | nABV | ND | ND | ND | 237 |
|  | ERR3313867 | TTV | 0.95 | 490 | 978 | 978 |
|  |  | HSV-1 | 0.95 | 863 | 934 | 908 |
|  |  | MEV | ND | ND | 493 | 507 |
|  |  | nABV | ND | ND | ND | 263 |
| Gire et al.<br>(2014) | SRR1564804 | GB Virus C | 0.58 | 9 | 80 | 70 |
|  |  | Chlamydia psittaci | 0.49 | 149481 | 548958 | 32519 |
|  | SRR1564809 | Lassa fever virus | 0.79 | 25160 | 142083 | 226975 |
|  | SRR1564816 | GB Virus C | 0.55 | 8 | 39 | 41 |
|  | SRR1564818 | GB Virus C | 0.98 | 56648 | 881381 | 927265 |
|  | SRR1564821 | HIV-1 | 0.61 | 263 | 4428 | 4383 |
|  | SRR1564825 | Enterovirus A | 0.45 | 476 | 11400 | 12182 |
|  | SRR1564831 | HIV-1 | 0.97 | 5971 | 104395 | 102412 |
|  | SRR1564833 | GB Virus C | 0.99 | 6185 | 18945 | 524 |
|  | SRR1564857 | EBOV | 1 | 11572 | 11567 | 11075 |
|  | SRR1972948 | EBOV | 1 | 18719 | 23447 | 18728 |
|  |  | Human<br>mastadenovirus C | 0.45 | 81 | 80 | 59 |
|  | SRR1972951 | EBOV | 1 | 487058 | 454225 | 432267 |
|  |  | Pseudomonas<br>aeruginosa | 0.49 | 1583 | 6790 | 16799 |
|  |  | Human<br>mastadenovirus C | 0.48 | 512 | 234 | 172 |
|  | SRR1972955 | EBOV | 1 | 75569 | 65403 | 63417 |
|  | SRR1972966 | EBOV | 1 | 790145 | 806519 | 733442 |
|  |  | Streptococcus pyogenes | 0.95 | 40356 | 71796 | 50841 |
|  | SRR1972969 | EBOV | 1 | 23116 | 14478 | 13906 |
|  |  | HIV-1 | 0.53 | 6 | ND | ND |
|  |  | Human mastadenovirus C | 0.41 | 111 | 44 | 37 |
|  | SRR1972971 | EBOV | 0.99 | 3904 | 3590 | 3277 |
|  |  | HIV-1 | 0.49 | 5 | 3 | 4 |

CZID similarly failed to identify nABV using nucleotide alignment. The pipeline was, however, able to assign 307 reads to *Orthobornavirus alphapsittaciforme*. While not strictly identifying this as a novel pathogen, the discrepancy between NT and NR results may prompt further investigations from an end-user. TaxTriage contains an optional DIAMOND module, which can be used to assess amino acid similarity if samples are suspected to contain a genetically divergent pathogen. Performance comparisons between TaxTriage and CZID were limited to nucleic acid similarity to evaluate default parameters in TaxTriage v2.0.8. Four of the eleven custom pipelines described in the COMPARE publication detected nABV (participants 3, 5, 6, and 12), although their analyses required a minimum of 12 and maximum of 60 hours to perform. These are compared to ∼2.5 hours from submission to result for CZID and only 15 minutes on a Mac OSX laptop with 32 GB RAM and an M2 Max processor for TaxTriage.

In addition, a single run (ERR3313866) from the COMPARE study failed to map any TTV reads, compared to 939 reads for both NT and NR for CZID. Upon investigation, ERR3313866 and ERR3313867 downloaded different reference sequences for TTV for minimap2 alignment. From the Kraken2 classification step, ERR3313866 reads were classified to taxID 68887 (Species = torque teno virus) while ERR3313867 reads were additionally classified to taxIDs 68887 and 3048427 (Species = Alphatorquevirus homin29). The Kraken2 results guided the download of reference sequences NC_076138 and NC_038344 for ERR3313866 and ERR3313867, respectively. The two reference sequences only share 56.1% identity by the Needleman-Wunsch algorithm and the COMPARE study authors almost certainly used a sequence more similar to NC_038344 to generate their *in silico* sequence. This may represent a disconnect between the reference sequences used to build the PlusPF database and those currently listed on RefSeq.

For the clinical pathogen detection challenge (CAMI), TaxTriage readily identified CCHFV with a TASS score of 0.50. In addition, TaxTriage identified a Human Immunodeficiency Virus-1 (HIV-1) coinfection (0.51 TASS score). CZID similarly detected CCHFV and HIV-1, although it is noted that CCHFV (*Orthonairovirus haemorrhagiae*) was not flagged as a known pathogen in the web GUI or the exported report. Upon further investigation, a mismatch between the TaxID listed for CCHFV on the CZID pathogens list (1980519) and the most current TaxID in NCBI (3052518) was noted. This discrepancy was reported to CZID using their contact form but has not, as of this writing, been corrected. Four of 10 challenge participants identified CCHFV in the sample (only three of which correctly diagnosed it as the causative agent of pathology), suggesting superiority of end-to-end pipelines over bespoke analysis approaches.

Lastly, TaxTriage results on the *Gire et al. (2014)* dataset demonstrate the clear value of metagenomics in an outbreak setting. Of the 20 suspected EVD patients, only seven were positive for Zaire Ebolavirus (EBOV) by mNGS. Four were HIV-1 positive (including two coinfected with EBOV), four were infected with GB Virus C (as reported elsewhere), and one was positive for Lassa Fever virus, which may present with similar symptomatology as EBOV as noted in *Gire et al. (2014)* ^50^. Additionally, TaxTriage provides some evidence for three separate cases of sepsis caused by *Chlamydia psittaci* (causative agent of Psittacosis), *Pseudomonas aeruginosa*, and *Streptococcus pyogenes*, none of which were reported in the original outbreak publication. CZID similarly detected all these coinfections, but at the cost of significantly less specificity. While the results in Table 4 show all primary pathogens with TASS Scores > 0.4, CZID reports nearly two magnitudes greater number of pathogens across most samples (Table S1). Identifying these agents takes a significant amount of manual curation when using CZID. These differences in curation are logical as TaxTriage is intentionally designed for high-confidence pathogen identification while CZID takes a much broader approach akin to community profiling.

In Table 4, COMPARE and CAMI datasets only include target pathogens. For *Gire et al. (2014)*, all pathogens with TASS Scores >= 0.4 are included. ND = Not Detected. MapQ scores are set at a minimum of 20 (default) for filtering purposes.

Interrogation of the potential pathogens identified by TaxTriage highlight several of the challenges with mNGS. The opportunistic pathogen with the highest TASS score (0.9) was *Staphylococcus cohnii* from the SRA sample SRR1564806. In addition, 13 of the 20 patients had detectable levels of *Pseudomonas fluoroscens*, with scores ranging from 0.44 to 0.61. Both of these species are commonly present in the skin microbiota ^51,52^, and thus are more likely to have been introduced during sample collection or processing than to be genuine causes of septicemia.

## Discussion

Rapid identification of emerging or unknown pathogens is an essential component of agile public health response. Within the United States and globally, many public health and research hospitals maintain capabilities of performing metagenomic or metatranscriptomic NGS. The ability to leverage such technologies for large-scale pathogen identification networks has so far been limited by challenges in scaling bioinformatics capacity across diverse local computational infrastructures and data-sharing restrictions, ultimately limiting the development of a coordinated monitoring capability.

The TaxTriage analytical pipeline has been developed to incorporate best-in-class metagenomic data analysis tools for high-confidence pathogen detection. Input from public health professionals, academic partners, and those working at the frontlines of unknown pathogen detection have guided development priorities ranging from installation requirements to data output formatting. In many instances, these developments have been guided by the operational needs of the public health response community. For example, in September 2023, TaxTriage was used to train a group of international medical professionals on advanced bioinformatics approaches for the detection of unknown pathogens, with the pipeline output used to evaluate annotation, alignment statistics, and data quality control. Within two days, attendees evaluated untargeted sequencing data to identify likely pathogens in the dataset, with further discussions focusing on potential reflex responses should a pathogen be identified. The feedback received at such sessions has guided features of the pipeline, including organisms to be included in the pathogens sheet, documentation for launching jobs via the command line, and design and structure of the Organism Discovery Report.

A remaining challenge to widespread adoption of NGS in clinical laboratories is the reproducibility of bioinformatic analyses. TaxTriage provides a consistent set and order of utilities to convert raw sequencing reads into a list of detected organisms and has been designed to maximize reproducibility. For example, the TaxTriage Organism Discovery Report incorporates results of reference alignment, rather than exact-match classification, as the default classifier (Kraken2) uses a probabilistic (and not deterministic) approach to assigning kmers to lowest common ancestors (LCAs) ^28^. While greatly reducing memory consumption compared to Kraken, the variability introduced by the minimizer approach in Kraken2 precludes the use of quantitative thresholds being applied to Kraken2 outputs for the purposes of clinical diagnostics. Use of deterministic methods for classification may increase reproducibility, but only if a consistent reference database is used^53^. Alignment-based approaches, as leveraged by TaxTriage, should ultimately be more reproducible due to the consistent approach to searching for sequence matches against an authoritative reference database, but it is important to note that the somewhat dynamic nature of the RefSeq representative assemblies (where new assemblies can be added or removed daily) can result in identical runs of TaxTriage spaced over time producing different quantitative results if the assembly for a classified species substantially changes. To overcome this, a local reference assembly can be used in place of dynamic retrieval of reference sequences from RefSeq.

In addition, taxonomic identification continues to be susceptible to errors and inconsistencies within reference databases. For example, initial analyses of the COMPARE dataset with early versions of TaxTriage completely failed to identify TTV. This failure was due to the provisional RefSeq sequence being downloaded for TTV (NC_076168) being derived from TTV-like mini virus with incorrect metadata. There may also be instances of errors in the generation of the sequence, as seen with the presence of parasitic RNA viruses in the assembly for *Toxoplasma gondii* ^54^. To reduce these issues, GenBank accessions were incorporated into TaxTriage as of v2.0.8.

While TaxTriage development has prioritized identification of human pathogens, the pipeline is highly amenable to use for agricultural or environmental pathogen detection. The host DNA removal utility within TaxTriage removes reads based on minimap2 alignment prior to classification. While the default option is the human assembly GRCh37, users can choose to select a genome from the illumina iGenomes dataset (inclusive of reference genomes for diverse species including cow, dog, horse, rat, and pig, among others), or upload a custom FASTA file to target specific host data. In addition, while the pathogen list was initially developed with a focus on human pathogens, its customizable nature enables users to ensure reference genomes for known pathogens of interest are included within the framework. The current version (as of date of publication) includes several known veterinary pathogens, based on the veterinary diagnostic test catalog of a collaborating institution.

Several technical considerations influence the operational deployment of TaxTriage. The requirement for identification via both a classification algorithm and nucleotide alignment to a classification-curated database increases the specificity but lowers the sensitivity for most organisms (except for organisms in the pathogens sheet which can bypass classification) when compared to approaches that require only classification or alignment and use much more comprehensive (yet computationally expensive) alignment databases (e.g., SURPI^55^). While this approach increases confidence in identified organisms, and therefore decreases costs and time associated with unnecessary reflex testing, it does increase the likelihood that some pathogens may fail to be identified. Additionally, as with nearly all automated pipelines, TaxTriage’s dependence on public reference sequences limits its ability to detect distinct or novel pathogens, as seen with the COMPARE dataset.

Metagenomic next-generation sequencing (mNGS) has been implemented as a CLIA-compliant laboratory-developed test (LDT) in specific, limited-use cases—primarily for sample types with low microbial background, such as cerebrospinal fluid and plasma. In these settings, test interpretation typically requires review by multiple qualified clinical microbiologists to ensure analytical validity and clinical relevance ^56,57^. Cross-sectoral activities are advancing individual components to facilitate broader leveraging of mNGS in clinical cases, including the development of wet– and dry-laboratory standards, electronic reporting of multiple potential pathogens, and discussions surrounding a regulatory roadmap for FDA clearance. Until these uncertainties are resolved, the use of mNGS to directly inform patient care will likely remain focused in specialized hubs. There is still substantial benefit for clinical laboratories or public health laboratories to invest in mNGS sequencing and analytics to inform outbreak investigations and intelligently direct reflexive testing. Public health laboratories can use mNGS as an agnostic screening method rather than a “test of last resort”. For samples for which standard assays for expected targets return negative results, mNGS can intelligently guide the use of follow-on, validated diagnostic tests, ultimately identifying putative pathogens for a greater proportion of samples, reducing the number of assays to be run, and lowering the per-sample cost of going from sample to answer. mNGS also has the added benefit of enabling pathogen genomics studies and identifying variants or clades, a level of resolution typically not afforded by assays such as quantitative polymerase chain reaction (qPCR).

TaxTriage will continue to be developed as OSS with planned activities including increasing the interpretability of the alignment confidence metric, expanding the number of classifiers and alignment algorithms available, and adding quality-of-life features to the Organism Discovery Report. Those interested in utilizing TaxTriage are encouraged to visit to the GitHub repository (https://github.com/jhuapl-bio/taxtriage/tree/main), which is released under the MIT License. The repository contains extensive documentation, example datasets, and video tutorials for launching and utilizing the pipeline in a variety of deployment settings. Bugs and feature requests can be indicated to the development team via the GitHub “Issues” feature.

## Acknowledgements

The authors acknowledge Robert Player and Hannah Martinez for their contributions to the TaxTriage codebase. The authors thank Tyler Wolford, Jarad Schiffer, and Christin Hanigan for supporting development of TaxTriage as well as several cross-sector partners that provided feedback on TaxTriage through deep dive working sessions. TaxTriage development was supported by the Cooperative Agreement Number NU60OE000104, funded by the Centers for Disease Control and Prevention through the Association of Public Health Laboratories. Implementation and training activities leveraging TaxTriage were supported by NIH Fogarty International Center under NAVSEA IDIQ Contract N00024-22-D-6404. The contents of this manuscript are solely the responsibility of the authors and do not necessarily represent the official views of the Centers for Disease Control and Prevention, the Department of Health and Human Services, or the Association of Public Health Laboratories.

## Supplementary Material

### Supplementary Methods

#### Confidence Score Derivation

To derive the TASS Score, TaxTriage uses the Gini Coefficient to measure the equality of coverage. While traditionally used for inequality analyses amongst economic environs, success has been identified using this metric for rapid analysis of variations in gene expression ^58^. A description of the Gini Coefficient, as applied to coverage distribution inequality, is shown in **Equations 1-3**. Our metric is an inverse of the traditional Gini Coefficient converted to the Average Fair Distribution Score (AFDS); for our reports, a coefficient of one describes perfect equality (and, thus, higher alignment confidence) while a coefficient of zero describes high levels of inequality (low alignment confidence). Efforts are underway to better define interpretability for the quantitative AFDS output scores. Additionally, we leverage the breadth of coverage and the drop in likely false positives when generating the *S*_1,02_from Sourmash-defined signatures using Sourmash version 4.18.14 **(Equation 4**). Finally, the aggregate of all three scores are calculated in **Equation 5**.

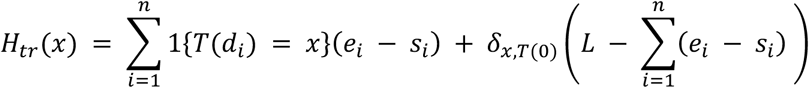

1. Compute the histogram (which is the distribution of depths and coverages for each reference
2. Calculate the G (Gini coefficient) from the depth profiles
3. Determine a scaling factor based on the genome length:
4. Compute dispersion factor:
5. Multiply these together to get the AFDS (Average Fair Distribution Score)

*δ_xt(0)_ is the is equal to 1 for any depth == *0* and *0* for empty positions (no coverage) 1 {T(d_i_) = x}* is equal to 1 if the depth of the current positions is the same as the last.

***Equation 1 – Representation of the alignment distribution confidence metric of depths and coverages for each species or strain***. *Reads must pass pre-filtering processes through fastp and the minimum mapq score before the confidence metric is calculated. The depth profiles are first converted into a histogram (H_tr_)*

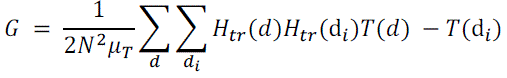

***Equation 2 – Representation of the alignment distribution confidence metric of depths and coverages for each species or strain****. The inequality factor is measured using a log-transformed inverse gini coefficient based on the depths of all positions. Each reference alignment profile is transformed into a histogram prior and the inequality of regions based on the depth and gaps is calculated*.

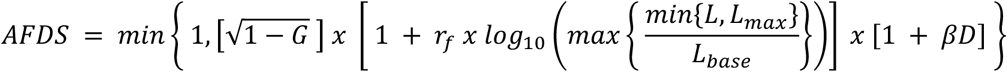

Where

● G be the raw Gini coefficient computed from the transformed coverage histogram *H_tr_*
● L be the gnome length
● *L_max_ is the full length of the genome or max it can be of all depths*
● *L_base_ is the baseline value*
● *r_f_ is the reward factor*
● α is the transformative partner
● β is the weight of the dispersion factor
● D is the positional factor

***Equation 3 – Representation of the alignment distribution confidence metric of depths and coverages for each species or strain****. Reads must pass pre-filtering processes through fastp and the minimum mapq score before the confidence metric is calculated. The depth profiles are first converted into a histogram (H_tr_)*.

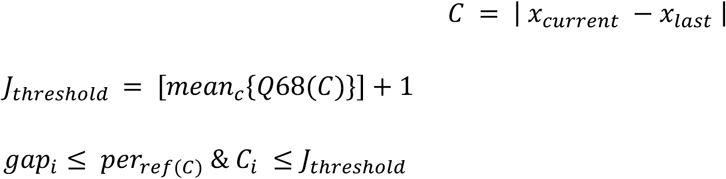

Where:

– C is the change in depth at each positional shift across the entire depth of coverage profiles for all references
– *J_threshold_* is the threshold determined by taking the mean of all changes and determining the 68th percentile + 1 depth. In short, changes in positions are then passed to the next step to be binned if the condition of the current depth x position is greater than the threshold
– A position is then merged IF both the previous step and the 0-depth length (from positions *x_i_* – *x_n_*) is greater than the defined threshold. The default width threshold is 3% of the breadth of the genome

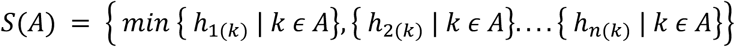

Where

– S(A) is the Minhash Signature of a set of reads binned to a single region
– *h_i_*_(*k*)_ is the hash value of the k-mer belonging to the region/subsequence from position 1 to n

The individual sets are compared as:

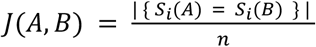

Where

– *S_i_*(*A*) and *S_i_*(*B*) are two distinct signature sets compared across references A and B
– n is the total number of hashes in total in the entire set of references.
– Any set of signatures that receive a match are then mapped to the original reads belonging to those regions. These reads are proportionately removed from the final alignment information based on the depth of coverage profiles to insulate against highly conserved regions in some species or genera.

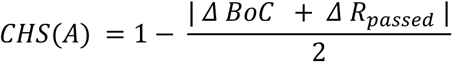

Where

– *CHS*(*A*) is the final Comparison Hash Score based on the change in both breadth of coverage and the drop in proportion of reads (equally divided) for any given reference A.

***Equation 4 – Determining the Signature Comparison Removals***. *Each alignment is binned based on a variety of regional characteristics using depth profile spikes/drops and empty regions of coverage. Regions are then binned and transformed into MinHash sketches using Sourmash. Similarities amongst regions are then identified and reads belonging to these portions are proportionally removed based on the total depth of coverage for each reference aligned. A final score is assigned based on the combination of both the* Δ *number of reads as well as the breadth of coverage*.

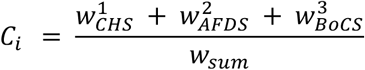

Where

– 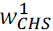 is the weighted form of *S*_1,02_
– 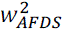 is the weighted form of the Average Fair Distribution Score
– 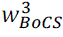 is the weighted Breadth of Coverage score

***Equation 5 – Representation of the final confidence metric score*.**

## Supplementary Tables and Figures

**Figure S1:**
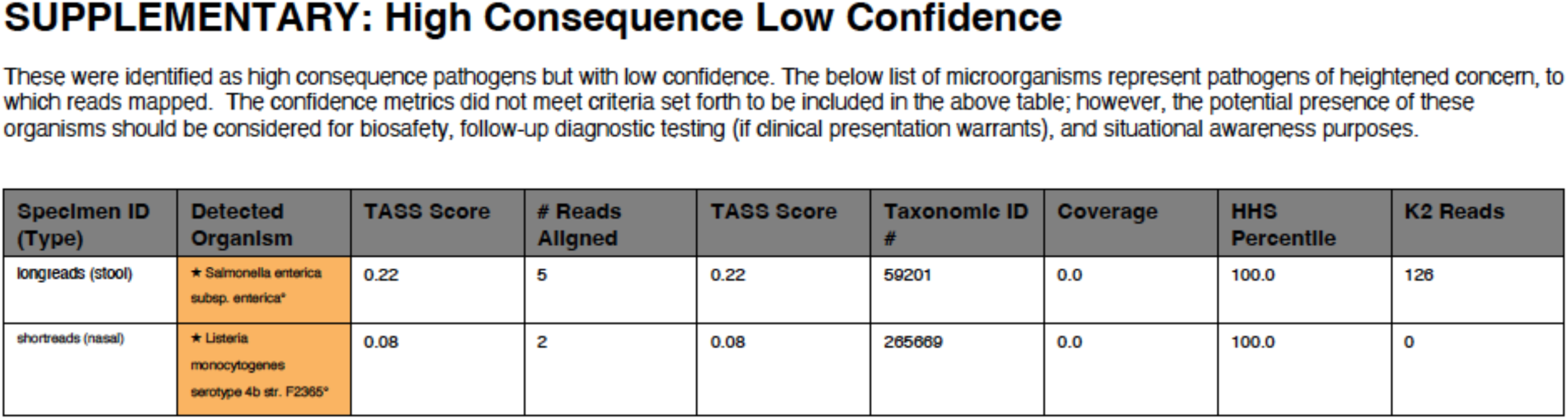
High Consequence Low Confidence Pathogens. Organisms annotated within the NIAID Biodefense Pathogens List are included in a separate supplementary table in the Organism Discovery Report if the TASS Score is <0.4. Although the confidence level of detection for these microorganisms is low, there are significant implications if truly present. This supplementary table is included for transparency to allow end user interpretation and potentially inform follow-on investigation if warranted.

**Figure S2:**
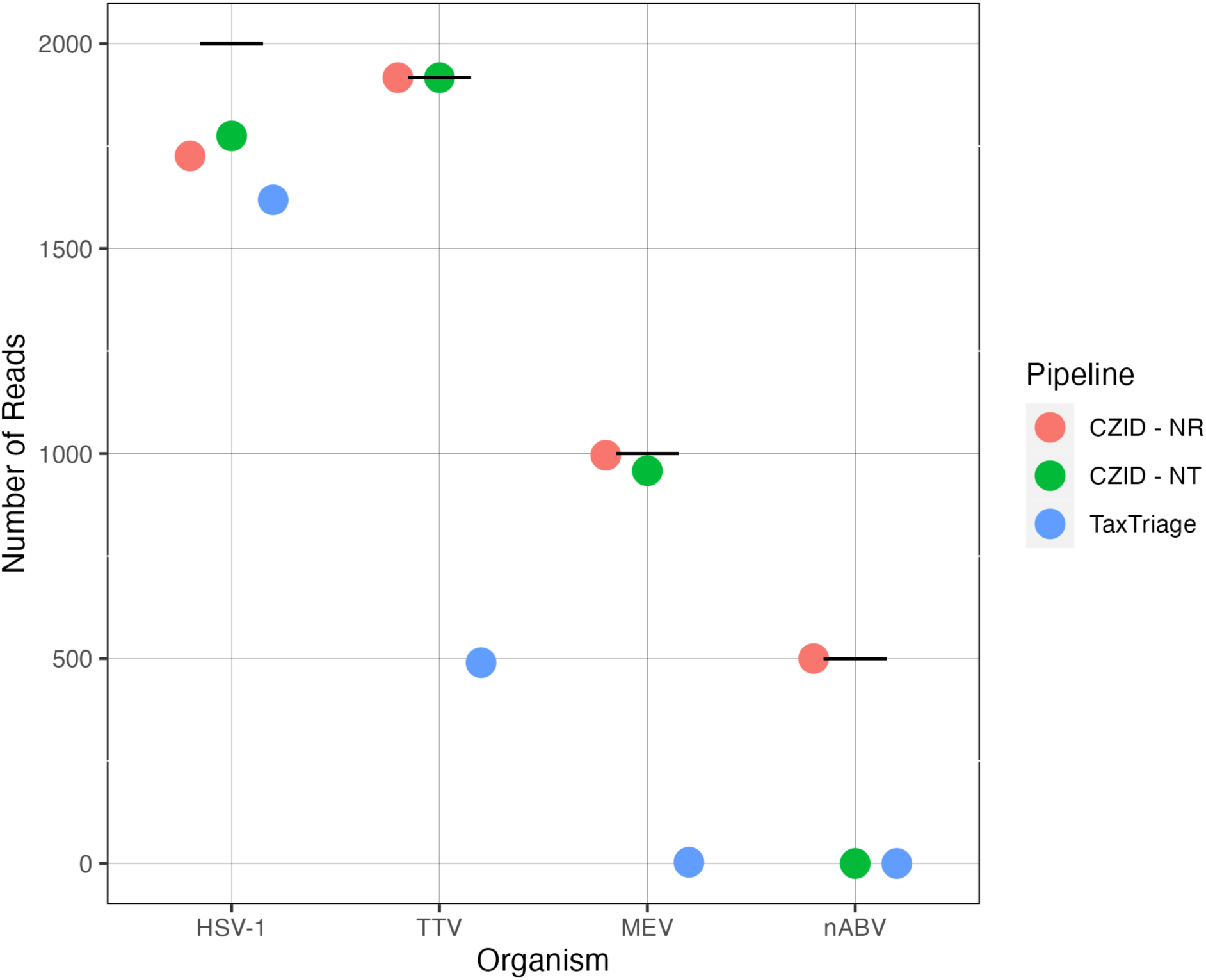
Read Recovery Rates for COMPARE dataset. Horizontal bar indicates ground truth number of synthetic reads summed across ERR3313866 and ERR3313867.

**Table S1.**
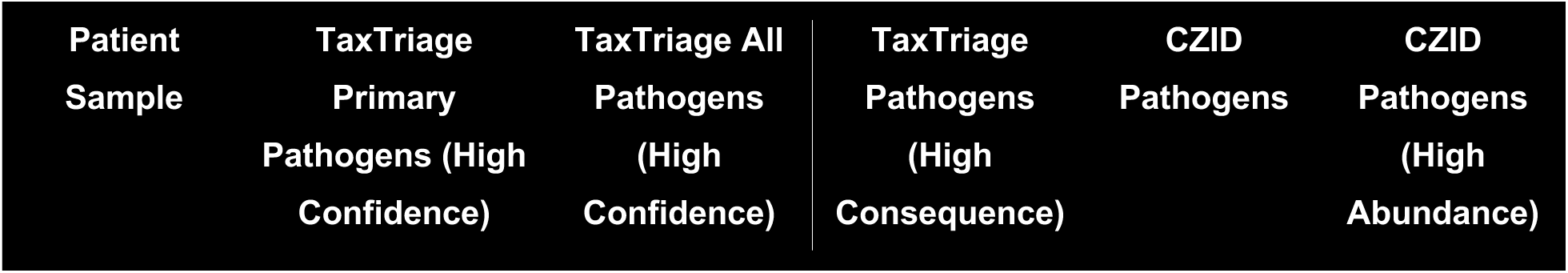
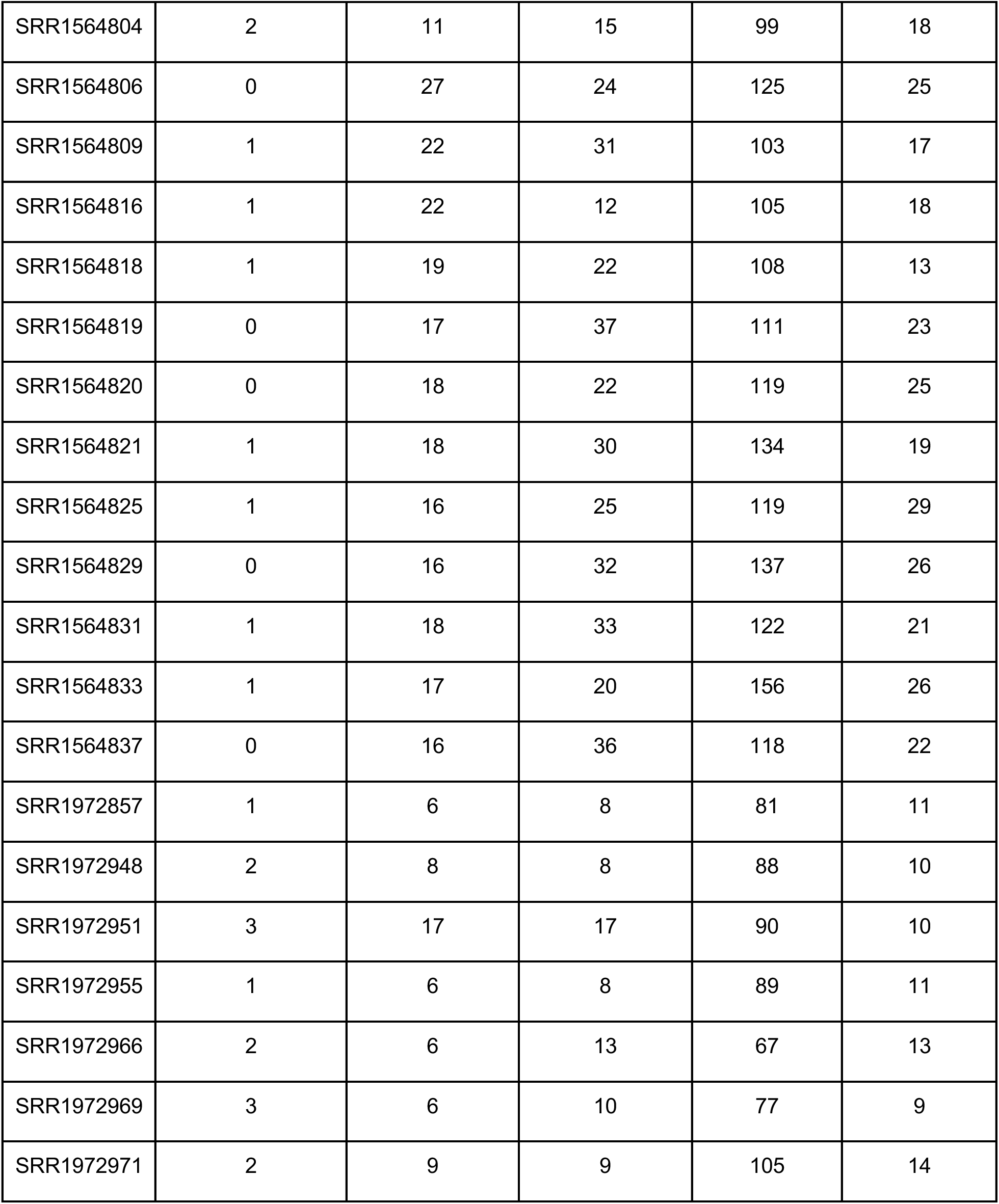
Total Pathogens identified in *Gire et al. (2014)* Table S1 depicts total pathogens identified by TaxTriage and CZID using data from *Gire et al. (2014)*. High confidence detections by TaxTriage are defined as organisms annotated as “Primary” pathogens with a TASS Score >= 0.4. All pathogens include organisms annotated as “Primary”, “Opportunistic”, or “Potential” with a TASS Score >= 0.4. High consequence detections by TaxTriage are defined as organisms annotated as “high consequence” pathogens regardless of TASS Score. Detections by CZID are defined as species-level organisms with a “known_pathogen” value of 1 with NT reads per million (rPM) > 1 and NR rPM > 1. High abundance detections by CZID are defined as species-level organisms with a “known_pathogen” value of 1 with NT reads per million (rPM) > 100 and NR rPM > 100.

